# Stress reactivity elicits a tissue-specific reduction in telomere length in ageing zebrafish (*Danio rerio*)

**DOI:** 10.1101/2020.04.17.046599

**Authors:** James R. Evans, Jose V. Torres-Pérez, Maria Elena Miletto Petrazzini, Riva Riley, Caroline H. Brennan

## Abstract

Telomere length reflects cellular ageing. Increased telomere shortening in leukocytes is associated with a range of neurodegenerative and cardiovascular diseases, the onset and progression of which may be mediated by behavioural traits such as anxiety and stress reactivity. However, the effects of the hypothalamus-pituitary-adrenal axis stress response are shown to be tissue specific. As such, leukocyte telomere length may not give an accurate measure of the relationship between stress-reactivity and telomere length in disease relevant tissues. To test the hypothesis that stress-reactivity contributes to age-related telomere shortening in a tissue specific manner, we examined the correlation between telomere length in heart and brain tissue and stress-reactivity in a population of young (6-9 month) and ageing (18 month) zebrafish. Stress-reactivity was assessed by tank diving, a zebrafish version of the rodent open-field test, and through gene expression. Telomere length was assessed using quantitative polymerase chain reaction. We show that ageing zebrafish have shorter telomeres in both heart and brain. Telomere length is inversely related to stress-reactivity in heart but not brain of ageing individuals. These data support the hypotheses that an anxious predisposition contributes to telomere shortening in heart tissue, and by extension age-related heart disease, and that stress-reactivity contributes to age-related telomere shortening in a tissue-specific manner.

## INTRODUCTION

Telomeres, tandem repeat guanine-rich DNA sequences, are specialised protective caps located at the ends of eukaryotic chromosomes. Crucial for the maintenance of genomic stability, they protect against the attrition of genetic material (1, 2), shortening with each cell division in most somatic tissues due to incomplete chromosome replication (3–5). Over time, progressively shortened telomeres may lead to cell-cycle arrest (6), apoptosis (7) or senescence (8). As such, telomere length may be considered as both a ‘mitotic clock’, reflecting cellular ageing, and a mechanism through which ageing, and consequently age-related disease, occurs.

However, telomere length may also be influenced by a wide range of factors in addition to cell replication. Chronic activation of the hypothalamus-pituitary-adrenal (HPA) axis, the main effector of the stress response, is associated with telomere shortening (9–11). Although under basal conditions the HPA axis is crucial for normal physiological function, a dysfunctional or prolonged response may contribute to telomere shortening through a variety of mechanisms (12). Notably, the initiation and termination of the HPA axis is thought to be mediated by the relative balance of glucocorticoid receptor (GR) and mineralocorticoid receptor (MR), signifying the importance of the GR:MR ratio in maintaining HPA axis homeostasis and regulating the stress response (13).

Neurodegenerative disease and heart disease are two of the most prevalent groups of age-related diseases. However, the role of telomere length in these diseases remains to be fully elucidated. Shorter telomeres have been identified in both Alzheimer’s (14) and Parkinson’s disease (15) patients and have been found to accelerate both the onset and underlying pathology of neurodegeneration in mice models (16). Shorter telomere length is also associated with a higher risk of coronary heart disease (17), an association that is thought to be causal (18). Interestingly, the likelihood of onset and progression of age-related diseases is associated with an anxious personality (19–21) and HPA axis dysfunction (22, 23). It is therefore thought that an exacerbated stress response may contribute to telomere shortening, accelerated ageing and, in turn, age-related disease (24–28). Additionally, telomere shortening can influence replicative senescence indirectly via downstream regulators such as SMAD specific E3 ubiquitin-ligase 3 (Smurf2) (29, 30) thus telomere length might be mediating other physiological aspects associated with ageing.

Many clinical studies pointing to a relationship between telomere length and age-related disease measure telomere length from blood samples, reporting leukocyte telomere length as a proxy for the tissues relevant to the disease (14, 15, 17). The true meaning of such research relies upon the fundamental assumption that changes in telomere length are robust across tissues and that leukocyte telomere length accurately reflects the tissue of interest, such as heart or brain. In support of this assumption, telomere shortening has been reported to be consistent across tissues irrespective of the proliferation rate of the tissue (4). In addition, although the enzyme telomerase is capable of elongating telomeres, and is expressed in stem cell compartments (31), it is postnatally supressed across all somatic tissues (3). However, other factors contributing to telomere dynamics may elicit tissue-specific effects. Of central importance to the current study is that both the effects of the HPA axis stress response (32, 33) and oxidative stress (34, 35) have been shown to operate in a tissue-specific manner, the latter of which has also been found to have a tissue-specific effect on telomere length (36). Thus, leukocyte telomere length may not accurately reflect the relationship between stress-reactivity and telomere length in different tissues relevant to disease.

Here, we used a population of zebrafish and a pseudo-longitudinal design to test the hypothesis that stress reactivity contributes to age-dependent telomere shortening in disease relevant tissues. Zebrafish have emerged as a promising animal model to explore telomere dynamics due to important similarities with humans. These include high genomic conservation (37), age-dependent telomere shortening (38–40), and, unlike mice, similar telomere lengths to humans (41, 42). Furthermore, the zebrafish hypothalamic-pituitary-interrenal (HPI) axis is analogous to its human counterpart, the HPA axis (43).

Differences in stress reactivity were determined by the novel tank paradigm in populations of young and ageing zebrafish (44). Fish occuring in the top and bottom of the population distribution curve for bottom dwelling were used to analysed heart and brain telomere length using qPCR. This technique was also used to assess relative expression of HPI axis related genes, including the glucocorticoid and mineralocorticoid receptors. We show that ageing zebrafish have shorter telomeres in both heart and brain tissue compared with young zebrafish. Furthermore, ageing zebrafish determined as high stress reactive have shorter telomeres in their hearts, but not brains, compared with the ageing fish with low stress reactivity. As such, this research demonstrates that stress reactivity may exacerbate age-related telomere shortening in a tissue-specific manner.

## RESULTS

### Selection of subjects for molecular analysis based on stress reactivity

In order to test the hypothesis that stress reactivity contributes to telomere shortening we first identified individual differences in the zebrafish anxiety-like behavioural response in populations of young (6-9 months) and ageing (18 month) zebrafish using the novel tank diving assay (Fig. 1.A).

**Figure 1.**
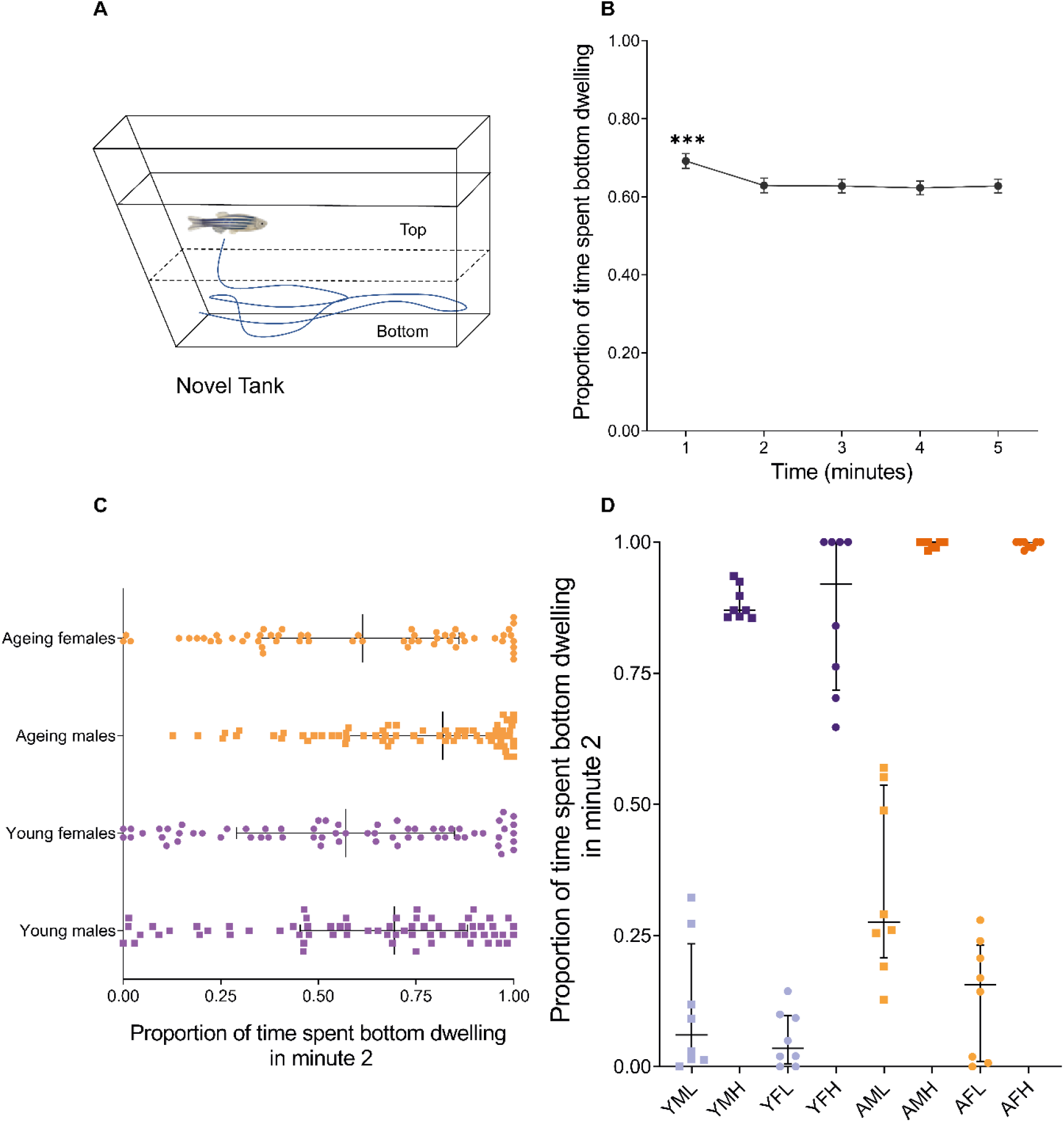
Zebrafish selected for molecular analysis based on stress reactivity. A) Diagram of the novel tank assay: each fish is individually transferred from its home tank to a novel test tank. The time (seconds) spent in the bottom in each minute was used for analysis. B) Entire population results for bottom dwelling by minute over the course of the novel tank test. C) Population distribution of young (6-9 month) and ageing (18 month) males and females based on their bottom dwelling in minute two (*N* = 256). D) Bottom dwelling differences in minute two of the fish selected for molecular analysis (*N* = 64). Note that D) corresponds to either extreme of the population distribution in C). All error bars represent ± standard error mean (SEM). YFL: young female low stress reactivity; YFH: young high stress reactivity; YML: young male low stress reactivity; YMH: young male high stress reactivity; AFL: ageing female low stress reactivity; AFH: ageing female high stress reactivity; AML: ageing male low stress reactivity; AMH: ageing male high stress reactivity. \*\*\**p*-value < 0.01

Overall, time was a significant predictor of bottom dwelling tendency (Likelihood Ratio Test (LRT) = 44.9, *p* < 0.001). In analysis of pairwise comparisons between bottom-dwelling tendency at each minute, the first minute was significantly different from all other timepoints (*p* <0.001 for all comparisons). The second minute was not significantly different than any subsequent timepoints (*p* > 0.98 for all comparisons, Fig. 1.B). Consequently, we chose the second minute to select individuals for molecular analysis (Fig. 1.C-D).

### Bottom dwelling tendency

The interaction between age and minute was a significant predictor of the proportion of time per minute spent dwelling on the bottom of the tank (LRT = 23.7, *p* < 0.001), however, the interaction between sex and minute was not significant (LRT = 1.9, *p* = 0.168). Sex was significant as a main effect (LRT = 8.2, *p* = 0.004), with males spending more time on the bottom (*M* = 0.688, *SD* = 0.258) than females (*M* = 0.583, *SD* = 0.315;). A post-hoc analysis revealed that ageing fish exhibited less bottom dwelling over time, but young fish did not change their bottom dwelling tendency over time (Fig. 2.A: see Supplementary Information for additional results)

**Figure 2.**
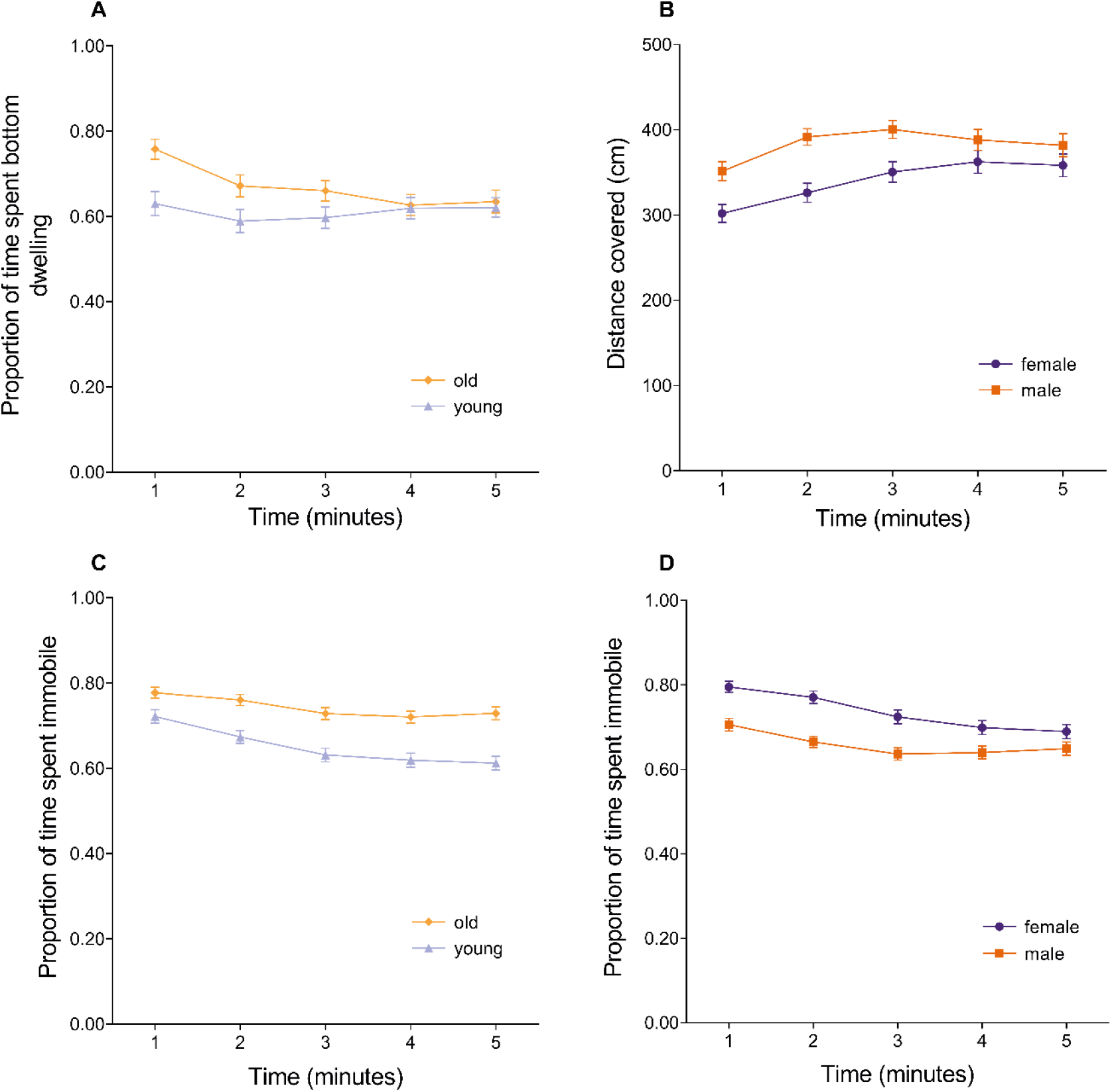
Results of the novel tank diving test. A) Overall proportion of time spent in the bottom third of the tank over the duration of the test. B) Distance covered by males and females for each minute of the novel tank test. C) Proportion of time ageing and young, and (D) males and females spent in an immobile state over the 5 minutes of the test. \*\**p*-value < 0.01

### Total distance moved

The interaction between sex and minute was a significant predictor of total distance moved per minute (LRT = 7.0, *p* = 0.008). However, the interaction between age and minute was not (LRT = 2.2, *p* = 0.137), nor was age a main effect (LRT = 2.2, *p* = 0.133). A post-hoc analysis revealed that males increased their distance moved at a faster rate than females (Fig. 2.B).

### Immobilization tendency

The interaction between age and minute was a significant predictor of the proportion of time per minute spent immobile (LRT = 6.5, *p* = 0.011), as was the interaction between sex and minute (LRT = 17.8, *p* < 0.001). Ageing fish spent more time immobile (proportion of time per minute spent immobile = 0.743, *SD* = 0.156) than young fish (proportion of time per minute spent immobile = 0.652, *SD* = 0.188; figure 2.C), and post-hoc analysis revealed that ageing fish resumed movement more slowly than younger fish. Females spent more time immobile (proportion of time per minute spent immobile = 0.736, *SD* = 0.177) than males (proportion of time per minute spent immobile = 0.659, *SD* = 0.173; Fig. 2.D), and post-hoc analysis revealed that females resumed movement more quickly than males.

### Ageing zebrafish with high stress reactivity have shorter telomeres in heart but not brain tissue

Telomere length shortened with age in both brain and heart irrespective of the level of stress reactivity (Fig. 3.A-B). Heart tissue of ageing subjects with high stress reactivity had significantly shorter telomeres compared with individuals with low stress reactivity (Fig. 3.A). No significant effect of stress-reactivity on telomere length in brain tissue was seen. No sex differences were identified in the telomere analysis.

**Figure 3.**
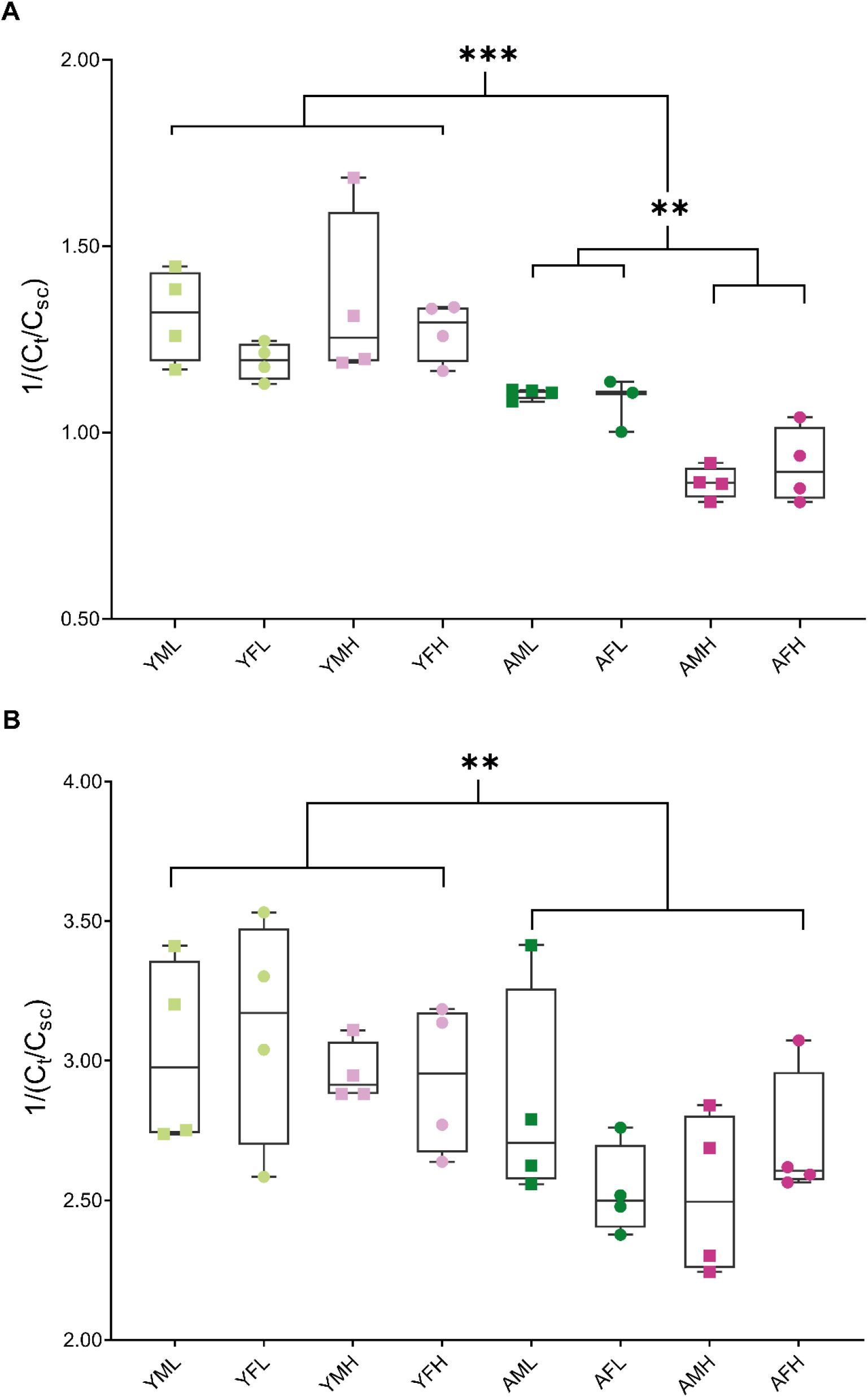
Telomere (TL) shortens with age and stress reactivity. A) Relative TL ratios calculated by qPCR for heart tissue grouped by age, sex and stress reactivity (*N* = 31). Age and stress reactivity in ageing zebrafish have an effect on TL. B) Relative TL ratios for brain grouped by age, sex and stress reactivity (*N* = 32). Only age has a significant effect. All error bars represent ± standard error mean (SEM). YFL: young female low stress reactivity; YFH: young high stress reactivity; YML: young male low stress reactivity; YMH: young male high stress reactivity; AFL: ageing female low stress reactivity; AFH: ageing female high stress reactivity; AML: ageing male low stress reactivity; AMH: ageing male high stress reactivity. \*\**p*-value < 0.01; \*\*\**p*-value < 0.01.

#### Heart Telomere Length

The three-way between-subjects ANOVA revealed that the main effect of age was significant (figure 3.A.; *F*(1,23) = 52.91, *p* < .001, partial η^2^ = .697). Shorter telomeres were found in ageing subjects (*M* = 0.98, *SD* = 0.12) compared with younger subjects (*M* = 1.28, *SD* = 0.14). The interaction between age and stress reactivity was significant (*F*(1,23) = 10.71, *p* = .003, partial η^2^ = .318). The Bonferroni adjusted post hoc comparisons found that ageing high stress reactive fish (*M* = 0.89, *SD* = 0.08) had significantly shorter telomeres in heart tissue compared with their low stress reactive siblings (*M* = 1.09, *SD* = 0.04) (figure 3.A; *p* = .002). At both extremes of the stress reactivity spectrum, ageing fish were found to have significantly shorter heart telomeres compared with their younger counterparts (young high stress reactive (*M* = 1.31, *SD* = 0.17) and young low stress reactive (*M* = 1.25, *SD* = 0.11), respectively: (*p* < .001; *p* = .011)). No other significant effects or interactions were found (Supplementary Tables 1 - 2).

#### Brain Telomere Length

The three-way between-subjects ANOVA revealed that the main effect of age was significant (figure 3.B: *F*(1,24) = 11.71, *p* = .002, partial η^2^ = .328). Shorter telomeres were found in ageing subjects (*M* = 2.65, *SD* = 0.29) compared with younger subjects (*M* = 3.01, *SD* = 0.28). No other significant effects or interactions were found (Supplementary Tables 1 - 2)

### HPI axis gene expression validates individual differences in behaviourally determined stress reactivity

Fish selected for telomere analysis were also tested for differences in HPI axis associated genes (Fig. 4.A-C). We performed qPCR on the expression of corticotropin-releasing factor, mineralocorticoid receptor and glucocorticoid receptor α. Subjects identified as more stress reactive in the novel tank assay were found to have significantly higher expression of corticotropin-releasing factor and mineralocorticoid receptor compared with their less stress reactive siblings; they were also found to have a significantly lower Mineralocorticoid:Glucocorticoid (MR:GR) ratio (Fig. 4.D).

**Figure 4.**
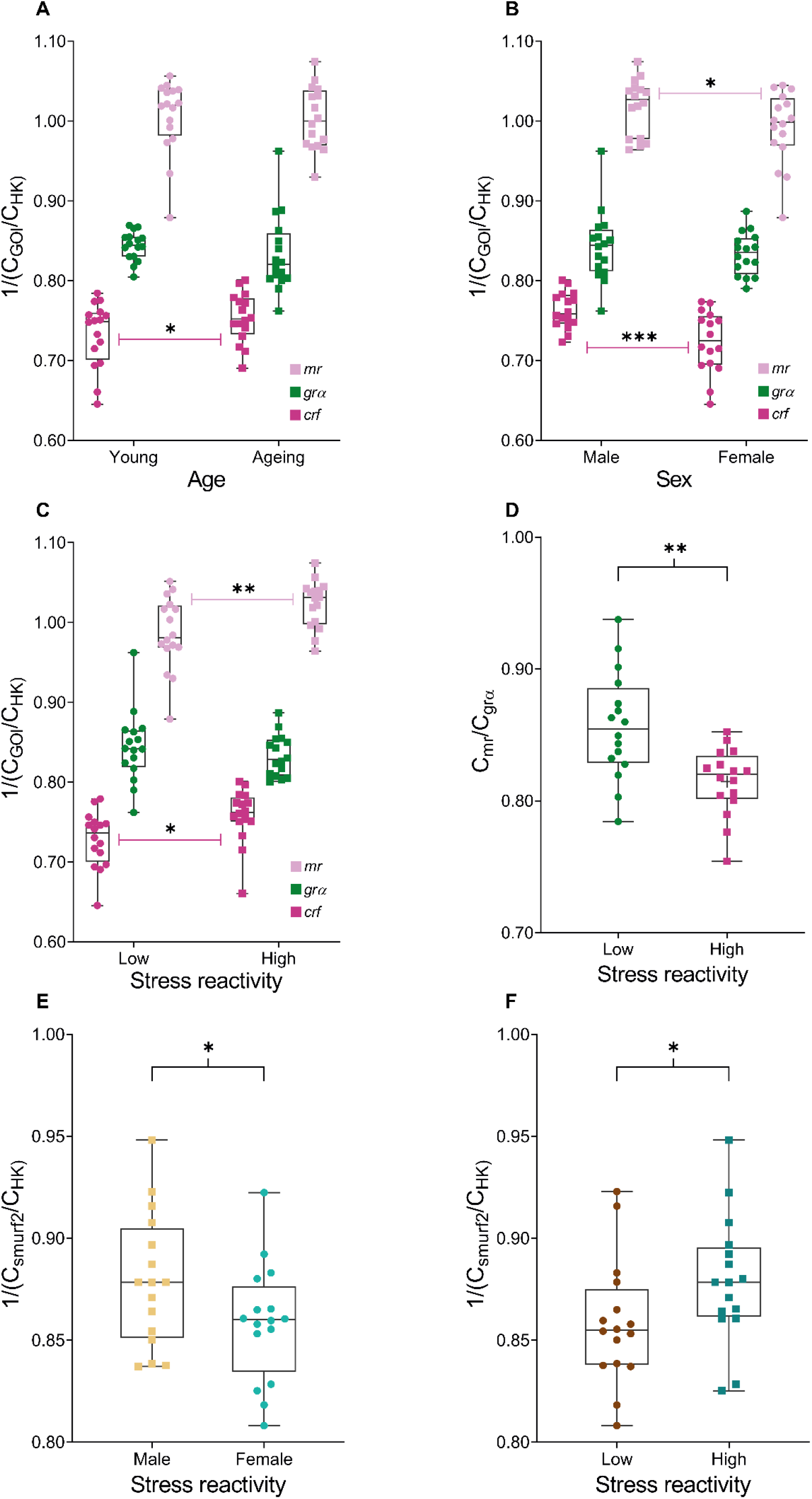
Levels of mRNA expression within brain extracts by qPCR validates behaviour in the novel tank. qPCR analysis of the HPI axis-associated genes, (corticotropin-releasing factor (crf), mineralocorticoid receptor (mr) and glucocorticoid receptor α (grα), in association with zebrafish A) age, (B) sex, and (C) stress reactivity (*N* = 96). D) Stress reactivity had a significant effect on the mr:grα ratio, a key indicator of HPA axis dysfunction (*N* = 32). The expression of SMAD Specific E3 Ubiquitin Protein Ligase (smurf2) was also analysed through qPCR and was significantly associated with both (E) sex and (F) stress reactivity (*N* = 32). *p-value < .005; \*\**p*-value < 0.01; \*\*\**p*-value < 0.01.

#### Corticotropin-releasing factor

The three-way between-subjects ANOVA revealed that the main effect of age was significant (*F*(1,24) = 4.80, *p* = .038, partial η^2^ = .167). Ageing subjects (*M* = 0.75, *SD* = 0.03) had significantly higher CRF expression compared with younger subjects (*M* = 0.73, *SD* = 0.04). The main effect of sex was significant (*F*(1,24) = 17.19, *p* < .001, partial η^2^ =.417). Males (*M* = 0.76, *SD* = 0.02) had significantly higher CRF expression compared with females (*M* = 0.72, *SD* = 0.04). The main effect of stress reactivity was also significant (*F*(1,24) = 9.99, *p* = .004, partial η^2^ = .294). Subjects with high stress reactivity (*M* = 0.76, *SD* = 0.03) had significantly higher CRF expression compared with low stress reactive subjects (*M* = 0.73, *SD* = 0.03).

#### Glucocorticoid receptor alpha

The three-way between-subjects ANOVA revealed no significant main effects nor any significant interactions (supplementary table 2).

#### Mineralocorticoid receptor (MR)

The three-way between-subjects ANOVA revealed that the main effect of sex was significant (*F*(1,24) = 0.04, *p* = .049, partial η^2^ = .002). Males (*M* = 1.02, *SD* = 0.04) had significantly higher MR expression compared to females (*M* = 0.99, *SD* = 0.05). The main effect of stress reactivity was also significant (*F*(1,24) = 4.31, *p* = .008, partial η^2^ = .152). High stress reactive subjects (*M* = 1.02, *SD* = 0.03) had significantly higher MR expression compared to subjects with low stress reactivity (*M* = 0.99, *SD* = 0.05).

#### Mineralocorticoid:Glucocorticoid Ratio

The three-way between-subjects ANOVA revealed that the main effect of stress reactivity was significant (*F*(1,24) = 12.17, *p* = .002, partial η^2^ = .336). Subjects with high behaviourally determined stress reactivity (*M* = 0.81, *SD* = 0.03) had a lower MR/GR ratio compared with low stress reactive zebrafish (*M* = 0.86, *SD* = 0.04).

### The SMAD specific E3 ubiquitin-ligase 3 (*smurf2*) is not a consequence of telomere attrition but is linked to stress reactivity

*smurf2* is thought to function downstream of telomere attrition (30). As such, we also tested whether the differences in telomere length attrition were accompanied by differences in the expression of *smurf2* by qPCR (figure 4.E-F). Pearson’s product-moment correlations were conducted to assess the relationship between telomere length, in both brain and heart, and *smurf2* gene expression. All variables were normally distributed, as assessed by Shapiro-Wilk’s test (*p* > .05). There was no statistically significant correlation between heart telomere length and *smurf2* expression (*r*(29) = −.163, *p* = .380), nor was there a statistically significant correlation between brain telomere length and *smurf2* expression (*r*(30) = −.025, *p* = .893). Nonetheless, the three-way between-subjects ANOVA revealed that the main effect of Sex was significant (*F*(1,24) = 4.35, *p* = .048, partial η^2^ = .153). Males (*M* = 0.88, *SD* = 0.03) had significantly higher *smurf2* expression compared to females (*M* = 0.86, *SD* = 0.03). The main effect of stress reactivity was also significant (*F*(1,24) = 4.35, *p* = .048, partial η^2^ = .153). Zebrafish with high stress reactivity (*M* = 0.88, *SD* = 0.03) had significantly higher *smurf2* expression compared with fish with low stress reactivity (*M* = 0.86, *SD* = 0.03).

## DISCUSSION

In the present study, we used zebrafish to test the hypothesis that stress reactivity exacerbates age-dependent telomere shortening and, importantly, that this occurs in a tissue-specific manner. From a population-size colony, fish with extreme high and low stress reactivity based on their performance in the novel-tank assay were selected for brain and heart TL analysis by qPCR. This procedure was carried out at two different stages of their mature life, young and ageing and their behaviourally determined stress reactivity was validated through the mRNA expression of HPI axis related genes. Increased age resulted in shorter TL in both organs. Strikingly, however, a higher anxious predisposition led to a significant shortening of telomeres in the hearts, but not the brains, of ageing fish. Our findings suggest that a high stress reactivity is a contributing factor for an increased rate of telomere attrition in zebrafish heart tissue and that stress-reactivity affects telomere length in a tissue-specific manner.

Zebrafish have been shown to demonstrate persistent individual differences in behaviour across both time and context. Thus, zebrafish provide a promising model for studying personality (45). Furthermore, individual differences in zebrafish anxiety-like behaviours have been shown to correspond with molecular differences, such as levels of dopamine and serotonin (46). Anxiety disorders are common in later life in humans (47, 48) and increased anxiety with ageing has been widely documented in rodents (49–51) and zebrafish (52). Our finding that ageing zebrafish exhibited increased anxiety-like behaviours (i.e. longer bottom dwelling and immobility state) compared with younger individuals is in line with such literature. It also correlate with the our reported changes in HPI related genes and the MR:GR ratio, thought to influence the initiation and termination of the stress response (44).

Furthermore, it has been shown that differences in behavioural response may be sexually dimorphic. Recently, Rambo and colleagues showed that females under stress had increased locomotor activity in comparison to males, although males had increased cortisol levels (53). Here, we also observed different stress reactivity in both sexes; males showed increased bottom-dwelling response, covered longer distances and were more mobile than females when exposed to a novel environment. Importantly, males also had significantly higher corticotropin-releasing factor, a key regulator of locomotor activity (54), and mineralocorticoid receptor expression, thus reflecting consistency between behavioural and molecular data.

Cardiovascular diseases have been widely associated with ageing, chronic stress and telomere shortening (55–57). The data reported here validates these associations and points to a possible mechanism: zebrafish with high stress reactivity have an increased or dysfunctional HPI axis response which, when chronically hyperactive, leads to an increased rate of telomere shortening in their hearts. The effect of stress related hormones such as cortisol and glucocorticoids on TL have been previously reported on peripheral tissues (55, 56, 58). The expression of telomerase, the enzyme with the ability to counteract telomere erosion, is susceptible to modulation by different hormones (11, 41, 58, 59), including glucocorticoids (60), thus pointing at a possible molecular mechanism.

Although zebrafish retain telomerase activity during their adult life (40, 42), our data shows a significant decline in TL with age, which may be further aggravated by HPI axis dysfunction. In most vertebrates including zebrafish, the major source of stem cells in the heart are the resident cardiomyocytes which can temporarily increase their telomerase activity (42, 57, 61, 62). When this peak is weakened by stress-related hormones (25, 41, 56) it may diminish their ability to counteract injuries thus leading to shorter telomeres and the associated increased risk of cardiomyopathies. However, our research cannot rule out a telomerase-independent process resulting from a chronically activated HPI axis, such as the postulated direct effect of reactive oxygen species (63). Nonetheless, the present investigation shows that zebrafish are a promising animal to model cardiomyopathies in ageing individuals.

A growing body of literature links short TL from peripheral proxy tissues, such as leukocytes, with the onset and progression of neurodegenerative and psychiatric disorders, and chronic stress (27, 55, 64–66). However, in the present study stress reactivity, although influential in heart telomere length, did not affect the rate of telomere attrition in the brain. Our finding that stress reactivity influences telomere length in a tissue-specific manner highlights the potential drawbacks of some clinical studies. Namely, that TL is either evaluated in superficial tissue for a short period of time or conducted post-mortem. Additionally, many of these studies are limited to relatively small samples in non-gender-balanced populations. Thus, the use of animal models is still a major need in ageing studies. The method described here uses a population-based zebrafish strategy to analyse TL in tissues relevant for ageing and disease (brain and heart) in a pseudo-longitudinal manner. If indeed anxiety predisposition does lead to exacerbated telomere shortening, this may only be observed in the brain at much older stages in zebrafish. Furthermore, differential rate of telomere shortening within different brain regions in humans has been previously described, with hippocampal differences significant between individuals with depression and controls (59, 67). Similarly, differences in the rate of telomere erosion have been observed within different cell types (68, 69). Therefore, tissue heterogeneity might be masking anxiety-driven differences within zebrafish brain extracts, a possibility that requires further investigation.

The upregulation of the ubiquitin ligase enzyme *smurf2* is suggested to be a consequence of telomere attrition in human fibroblasts (30). Its expression is proposed to be independent of the levels of cellular stress and DNA damage and though to induce senescence (29, 30). However, telomere shortening did not predict *smurf2* expression in our study. Instead, we found that its expression was increased in the anxious phenotype regardless of age. Thus, suggesting that the increase in *smurf2* is not a consequence of telomere shortening in zebrafish. In accordance with others (29), we observed a sexual dimorphic pattern with increased expression of *smurf2* in males.

Although we report TL shortening at 18 months of age in both brain and heart tissue, previous research using zebrafish has not observed age-dependent differences in TL until fish are at least 2 years of age (41, 70).Furthermore, others have shown no TL effect of age in the brain of zebrafish (39). Failure to detect telomere shortening in brain tissues could be a consequence of the lower rate of cell renewal and neurogenesis in the adult brain compared to heart making detection difficult (39, 63, 69).The lack of consistency in zebrafish findings highlights possible differences in the technique used to assess telomere length, on the rate of telomere attrition between strains (40, 41) and animal facilities, which should all be carefully considered. Furthermore, we have combined data from a sibling populations of young and ageing zebrafish, thus allowing for samples to be processed simultaneously, which assures similar storage time and reagents used, critical consideration for the study of telomere biology (71).

Taken together, we have demonstrated that an anxious predisposition, manifested as elevated stress reactivity, affects telomere length in zebrafish in a tissue-specific manner, exacerbating age-dependent telomere shortening in heart but not brain tissue. As such, we demonstrate the importance of assessing TL in disease-relevant tissue to obtain meaningful results. Furthermore, we have validated a population-base method using zebrafish that allows for pseudo-longitudinal analysis of telomere attrition in a tissue-specific manner.

## MATERIALS AND METHODS

### Subjects

256 adult Tuebingen (TU, wild type) zebrafish (*Danio rerio*), 134 males and 122 females, were reared to 6-9 months (133 subjects) or 18 months (123 subjects) of age in The Zebrafish Facility at Queen Mary University of London (QMUL). Fish were housed in a recirculating system (Techniplast, UK) with a light:dark cycle of 14:10 and a constant temperature of 28.4°C. All subjects were fed twice daily with ZM-400 fry food in the morning and brine shrimp in the afternoon. All animal procedures in this study were reviewed by the QMUL ethics committee (AWERB) and conducted in accordance with the Animals (Scientific Procedures) Act, 1986 and Home Office Licenses.

### Behavioural experiments

All fish were exposed to the novel tank diving test as described before (37). Briefly, after at least one hour of acclimation to the test room, zebrafish were individually tested in a 3.5-L trapezoid-shaped tank (17.5 cm height × 11.5 cm width × 28.5 cm top length × 24.5 cm bottom length) filled with system water (Fig. 1 A). Fish behaviour was recorded for 5 min from the side using a DMK 21AF04 Firewire Monochrome camera for five minutes. This period commenced immediately on their introduction to the new tank. Swimming activity, including the parameters of bottom dwelling (i.e. time spent in the lower third of the tank sectioned horizontally), mobility state (i.e. changes in the animal body displacement) and distance covered, were tracked and analysed using EthoVision software (Noldus Information Technology, Wageningen, NL). On completion of the assay the subjects were single housed until tissue harvesting. Water was replaced for each trial.

### Dissection

Fish used for dissection were selected based on bottom dwelling results (Fig. 1.B). Fish were sorted as high/low stress reactive based on their bottom dwelling time in minute 2 (Fig. 1 C). We selected eight fish from the top and the bottom of the distribution for each age and sex (N = 64) (Fig. 1 D). The fish were euthanised with tricaine methane sulfonate (MS222). Samples of whole-brain and heart tissue were dissected out, immediately snap-frozen and stored at −80°C until required.

### Extracting genomic DNA

Genomic DNA was isolated from whole-brain and heart samples using the HotSHOT protocol (64). In summary, 100 μL of 50 mM NaOH was added to each sample and then incubated at 95°C for 30 minutes in a heat block. Once completed, 10 μL of 1 M Tris-HCL, pH 8.0, was added to neutralise pH.

### Extracting mRNA and cDNA library generation

Total RNA was isolated from whole-brain samples using TRIzol™. The protocol followed is as stated in the TRIzol™ Reagent User Guide (Invitrogen). Briefly, following sample homogenisation with TRIzol™, RNA is isolated through precipitation, washing, and resuspension. RNA concentrations and quality were checked using a Thermo Scientific™ NanoDrop 2000. Corresponding cDNA libraries were generated using the ProtoScript® II First Strand cDNA Synthesis Kit (NEB (UK) Ltd., Hitchin, UK).

### Real-time quantitative PCR

All primer sequences can be found in Supplementary Table 1. In order to investigate differences in HPI axis and Smurf2 gene expression, relative qPCR assays were then performed using SYBR Green (Applied Biosystems) and the CFX Connect™ Real-Time System (Bio-Rad), with all reactions performed in triplicate. Reference genes for these qPCR analyses were β-actin and rpll3α. PCR reactions were performed at 95°C for 5 minutes followed by 50 cycles of 95°C for 10 seconds, 60°C for 12 seconds and 72°C for 12 seconds. Relative mRNA expression, gene quantification cycle (Cq) values were normalised against reference gene: 1/(Cq_GeneOfInterest_/Cq_houskeeping_).

To analyse relative average TL of the samples, relative qPCR was again conducted using SYBR Green (Applied Biosystems) and the CFX Connect™ Real-Time System (Bio-Rad), with all reactions performed in triplicate. The protocol used was a modified version of the method described by Cawthon (59). TL qPCR reactions were performed at 95°C for 10 minutes followed by 40 cycles of 95°C for 15 seconds and 54°C for 2 minutes. Relative TL was expressed as 1/(Cq_telomere_/Cq_SingleCopyGene_). The single copy gene used as reference for the relative ratio was cfos, primers without intra-exon introns (supplementary table 1). Missing data were assigned the highest measured Cq for that gene of interest plus one. Primers had similar efficiency values (Supplementary Table 3).

### Data analysis

All statistical behavioural analyses were carried out in R version 3.2.2 (R core developer team), and linear mixed effects models (LME) were fitted using the lme4 package (72); generalized mixed effect models (GLMM) were fitted using the glmmTMB package (73). Data distributions were initially assessed visually, and model diagnostics were subsequently checked to assure appropriate fits. We hence fitted LMEs for normally distributed data (total distance moved) and GLMMs with betabinomial error distributions where binomial models had been overdispersed (74). We used the emmeans package in R (Lenth, 2019) to perform Tukey posthoc tests. Unless indicated, the details of our posthoc analysis are contained in our supplementary materials.

To choose a timepoint at which to select individuals for our molecular analysis of telomere length, we performed a Tukey posthoc test on the bottom dwelling data pooled across age and sex to analyze which timepoint represented the earliest ‘recovery’ from the stress. We constructed a betabinomial GLMM based on this pooled data with the proportion of time spent swimming on the bottom as the response variable, minute as the fixed effect, and batch number and individual ID as random effects.

To assess the effect of age and sex on the tendency to swim on the bottom over the course of the experiment, we ran a betabinomial GLMM with the proportion of time spent swimming on the bottom in each minute as the response variable, the interaction between age and minute and the interaction between sex and minute as fixed effects, and individual ID and batch as random effects. Interactions that were not significant were subsequently refitted as main effects.

To assess the effect of age and sex on the change in distance swam per minute over the course of the experiment, we ran an LME with total distance moved per minute as the response variable, the interaction between age and minute and the interaction between sex and minute as fixed effects, and individual ID and batch as random effects. Interactions that were not significant were subsequently refitted as main effects.

To assess the effect of age and sex on the tendency to remain immobile over the course of the experiment, we ran a betabinomial GLMM with the proportion of time spent swimming on the bottom in each minute as the response variable, the interaction between age and minute and the interaction between sex and minute as fixed effects, and individual ID and batch as random effects.

Differences in gene expression and telomere length were assessed using three-way between-subjects ANOVAs with age (ageing, young), sex (male, female), and stress reactivity (low, high) as the independent variables. All follow-up tests generating simple effect comparisons were conducted with the use of SPSS syntax and to account for multiple testing Bonferroni corrections were applied. Pearson’s product-moment correlations were run to assess the relationship between telomere length and *smurf2* gene expression. All effects are reported as significant at p <.05. All molecular analyses were conducted using SPSS Statistics Version 26 (IBM Corporation, 2019).

## AKNOWLEDGEMENTS

Funders: Human Frontiers Scientific Program (RGP 0008/2017, CHB, JVTP), Leverhulme Trust. (RPG-2016-143, RR, CHB), BBSRC (BB/M007863, CHB). MEMP is funded by the European Union’s Horizon 2020 research and innovation programme (Marie Sklodowska-Curie Action 750200). CHB is a member of the Royal Society Industry Fellows’ College.

