## Supplementary materials for "Stress reactivity elicits a tissue-specific reduction in telomere length in ageing zebrafish (*Danio rerio*)"

**Supplementary Information for additional behavioural results:**

Bottom Dwelling

A post-hoc analysis revealed that ageing fish exhibited a significant decrease in bottom dwelling tendency from minute one to minutes two, three, four, and five ($p < 0.001$), while young fish did not show a change in bottom dwelling tendency from minute one to any other minute ($p = 0.247$; figure 2.A). No other comparisons between minutes two, three, four, or five were significant in either age group.

Total Distance Moved

A post-hoc analysis revealed that males exhibited a significant increase in distance moved from minute one to minute two ($p = 0.005$), while females did not show a significant increase in distance moved from minute one to minute two ($p = 0.452$); rather, females exhibited a significant increase in distance moved from minute one to minute three ($p < 0.001$) and all other minutes ($p < 0.001$).

Immobility

A post-hoc analysis revealed that ageing fish did not exhibit a change in immobilization from minute one to minute two ($p = 0.938$) but did exhibit a significant decrease in immobilization from minute one to all subsequent minutes ($p < 0.001$); no other comparisons were significant. Younger fish exhibited significantly decreased immobilization from minute one to minute two ($p = 0.003$) as well as between minutes one and two and all subsequent minutes (p < 0.040). In addition, a post-hoc test revealed that females first exhibited a significantly lower immobilization tendency between minute one and minute two ($p = 0.003$), whereas males did not exhibit a significant different between minute one and minute two ($p = 0.938$); rather, their immobilization tendencies was first significantly different between minute one and minute three ($p = 0.001$).

### Supplementary Table 1. Descriptive statistics for molecular data

**Main effects:**

| | Age | | | | Sex | | | | Stress reactivity | | | |
|---|---|---|---|---|---|---|---|---|---|---|---|---|
| | Old | | Young | | Male | | Female | | High | | Low | |
| | *M* | *SD* | *M* | *SD* | *M* | *SD* | *M* | *SD* | *M* | *SD* | *M* | *SD* |
| Heart Telomere Length | 0.98 | 0.12 | 1.28 | 0.14 | 1.16 | 0.23 | 1.12 | 0.16 | 1.10 | 0.25 | 1.18 | 0.12 |
| Brain Telomere Length | 2.65 | 0.29 | 3.01 | 0.28 | 2.84 | 0.34 | 2.82 | 0.34 | 2.78 | 0.28 | 2.88 | 0.38 |
| Corticotropin-releasing factor | 0.75 | 0.03 | 0.73 | 0.04 | 0.76 | 0.02 | 0.72 | 0.04 | 0.76 | 0.03 | 0.73 | 0.03 |
| Glucocorticoid receptor alpha | 0.83 | 0.05 | 0.84 | 0.02 | 0.84 | 0.04 | 0.83 | 0.03 | 0.83 | 0.03 | 0.84 | 0.04 |
| Mineralocorticoid receptor | 1.00 | 0.04 | 1.01 | 0.05 | 1.02 | 0.04 | 0.99 | 0.05 | 1.02 | 0.03 | 0.99 | 0.05 |
| Mineralocorticoid:Glucocorticoid Ratio | 0.83 | 0.04 | 0.84 | 0.04 | 0.83 | 0.03 | 0.84 | 0.04 | 0.81 | 0.03 | 0.86 | 0.04 |
| SMAD specific E3 ubiquitin-ligase 3 | 0.87 | 0.03 | 0.87 | 0.03 | 0.88 | 0.03 | 0.86 | 0.03 | 0.88 | 0.03 | 0.86 | 0.03 |

**Two-way interactions:**

**Age and sex**

| | Old | | | | Young | | | |
|---|---|---|---|---|---|---|---|---|
| | Male | | Female | | Male | | Female | |
| | *M* | *SD* | *M* | *SD* | *M* | *SD* | *M* | *SD* |
| Heart Telomere Length | 0.98 | 0.13 | 0.98 | 0.12 | 1.33 | 0.17 | 1.23 | 0.08 |
| Brain Telomere Length | 2.68 | 0.36 | 2.62 | 0.21 | 2.99 | 0.23 | 3.02 | 0.33 |
| Corticotropin-releasing factor | 0.77 | 0.03 | 0.74 | 0.03 | 0.76 | 0.02 | 0.71 | 0.04 |
| Glucocorticoid receptor alpha | 0.83 | 0.06 | 0.83 | 0.03 | 0.85 | 0.01 | 0.84 | 0.02 |
| Mineralocorticoid receptor | 1.01 | 0.04 | 1.00 | 0.04 | 1.03 | 0.02 | 0.98 | 0.05 |
| Mineralocorticoid:Glucocorticoid Ratio | 0.83 | 0.05 | 0.83 | 0.03 | 0.83 | 0.01 | 0.85 | 0.05 |
| SMAD specific E3 ubiquitin-ligase 3 | 0.87 | 0.03 | 0.86 | 0.04 | 0.88 | 0.03 | 0.85 | 0.02 |

**Age and stress reactivity**

| | Old | | | | Young | | | |
|---|---|---|---|---|---|---|---|---|
| | High | | Low | | High | | Low | |
| | *M* | *SD* | *M* | *SD* | *M* | *SD* | *M* | *SD* |
| Heart Telomere Length | 0.89 | 0.08 | 1.09 | 0.04 | 1.31 | 0.17 | 1.25 | 0.11 |
| Brain Telomere Length | 2.62 | 0.27 | 2.69 | 0.32 | 2.94 | 0.19 | 3.07 | 0.35 |
| Corticotropin-releasing factor | 0.77 | 0.02 | 0.73 | 0.03 | 0.74 | 0.04 | 0.72 | 0.04 |
| Glucocorticoid receptor alpha | 0.83 | 0.03 | 0.84 | 0.06 | 0.84 | 0.02 | 0.85 | 0.02 |
| Mineralocorticoid receptor | 1.02 | 0.04 | 0.99 | 0.04 | 1.03 | 0.02 | 0.99 | 0.06 |
| Mineralocorticoid:Glucocorticoid Ratio | 0.81 | 0.04 | 0.85 | 0.04 | 0.82 | 0.01 | 0.86 | 0.04 |
| SMAD specific E3 ubiquitin-ligase 3 | 0.89 | 0.02 | 0.85 | 0.03 | 0.87 | 0.04 | 0.87 | 0.02 |

**Sex and stress reactivity**

| | Male | | | | Female | | | |
|---|---|---|---|---|---|---|---|---|
| | High | | Low | | High | | Low | |
| | *M* | *SD* | *M* | *SD* | *M* | *SD* | *M* | *SD* |
| Heart Telomere Length | 1.11 | 0.30 | 1.21 | 0.14 | 1.09 | 0.21 | 1.14 | 0.08 |
| Brain Telomere Length | 2.74 | 0.31 | 2.94 | 0.35 | 2.82 | 0.26 | 2.82 | 0.42 |
| Corticotropin-releasing factor | 0.78 | 0.02 | 0.75 | 0.02 | 0.74 | 0.04 | 0.71 | 0.03 |
| Glucocorticoid receptor alpha | 0.83 | 0.02 | 0.85 | 0.06 | 0.83 | 0.03 | 0.83 | 0.03 |
| Mineralocorticoid receptor | 1.02 | 0.04 | 1.01 | 0.03 | 1.02 | 0.02 | 0.96 | 0.04 |

| | | | | | | | | |
|---|---|---|---|---|---|---|---|---|
| Mineralocorticoid:Glucocorticoid Ratio | 0.81 | 0.03 | 0.84 | 0.04 | 0.82 | 0.02 | 0.87 | 0.04 |
| SMAD specific E3 ubiquitin-ligase 3 | 0.89 | 0.03 | 0.87 | 0.04 | 0.87 | 0.03 | 0.85 | 0.02 |

**Three-way interaction:** **Age, sex and stress reactivity**

| | **Old** | | | | | | | |
|---|---|---|---|---|---|---|---|---|
| | **Male** | | | | **Female** | | | |
| | **High** | | **Low** | | **High** | | **Low** | |
| | ***M*** | ***SD*** | ***M*** | ***SD*** | ***M*** | ***SD*** | ***M*** | ***SD*** |
| Heart Telomere Length | 0.87 | 0.04 | 1.10 | 0.01 | 0.91 | 0.10 | 1.08 | 0.07 |
| Brain Telomere Length | 2.52 | 0.29 | 2.85 | 0.39 | 2.71 | 0.24 | 2.53 | 0.16 |
| Corticotropin-releasing factor | 0.78 | 0.02 | 0.75 | 0.02 | 0.77 | 0.01 | 0.72 | 0.02 |
| Glucocorticoid receptor alpha | 0.81 | 0.01 | 0.86 | 0.09 | 0.84 | 0.04 | 0.82 | 0.03 |
| Mineralocorticoid receptor | 1.01 | 0.05 | 1.00 | 0.04 | 1.03 | 0.02 | 0.97 | 0.03 |
| Mineralocorticoid:Glucocorticoid Ratio | 0.80 | 0.05 | 0.85 | 0.06 | 0.82 | 0.03 | 0.85 | 0.03 |
| SMAD specific E3 ubiquitin-ligase 3 | 0.89 | 0.02 | 0.86 | 0.04 | 0.98 | 0.02 | 0.84 | 0.03 |

| | **Young** | | | | | | | |
|---|---|---|---|---|---|---|---|---|
| | **Male** | | | | **Female** | | | |
| | **High** | | **Low** | | **High** | | **Low** | |
| | ***M*** | ***SD*** | ***M*** | ***SD*** | ***M*** | ***SD*** | ***M*** | ***SD*** |
| Heart Telomere Length | 1.35 | 0.23 | 1.31 | 0.12 | 1.27 | 0.08 | 1.19 | 0.05 |
| Brain Telomere Length | 2.95 | 0.11 | 3.03 | 0.34 | 2.93 | 0.27 | 3.11 | 0.41 |
| Corticotropin-releasing factor | 0.77 | 0.01 | 0.75 | 0.02 | 0.72 | 0.04 | 0.70 | 0.04 |
| Glucocorticoid receptor alpha | 0.85 | 0.02 | 0.85 | 0.02 | 0.83 | 0.02 | 0.84 | 0.02 |
| Mineralocorticoid receptor | 1.04 | 0.02 | 1.02 | 0.03 | 1.01 | 0.02 | 0.95 | 0.06 |
| Mineralocorticoid:Glucocorticoid Ratio | 0.82 | 0.00 | 0.83 | 0.01 | 0.81 | 0.02 | 0.89 | 0.05 |
| SMAD specific E3 ubiquitin-ligase 3 | 0.90 | 0.04 | 0.87 | 0.03 | 0.84 | 0.02 | 0.86 | 0.01 |

### Supplementary Table 2. ANOVA molecular data results for main effects and interactions

| **Main effects:** | **Age** | | | | **Sex** | | | | **Stress reactivity** | | | |
|---|---|---|---|---|---|---|---|---|---|---|---|---|
| | ***df*** | **F** | ***p*** | **$\eta_p2$** | ***df*** | **F** | ***p*** | **$\eta_p2$** | ***df*** | **F** | ***p*** | **$\eta_p2$** |
| Heart Telomere Length | **1,23** | **52.91** | **<.001*** ** | **0.697** | 1,23 | 1.17 | 0.292 | 0.048 | 1,23 | 3.46 | 0.076 | 0.131 |
| Brain Telomere Length | **1,24** | **11.71** | **0.002** ** | **0.328** | 1,24 | 0.02 | 0.900 | 0.001 | 1,24 | 0.94 | 0.342 | 0.038 |
| Corticotropin-releasing factor | **1,23** | **4.80** | **0.038*** | **0.167** | **1,23** | **17.19** | **<.001*** ** | **0.417** | **1,23** | **9.99** | **0.004** ** | **0.294** |
| Glucocorticoid receptor alpha | 1,23 | 0.58 | 0.453 | 0.024 | 1,23 | 0.52 | 0.480 | 0.021 | 1,23 | 0.67 | 0.420 | 0.027 |
| Mineralocorticoid receptor | 1,23 | 8.30 | 0.839 | 0.257 | **1,23** | **0.04** | **0.049*** | **0.002** | **1,23** | **4.31** | **0.008** ** | **0.152** |
| Mineralocorticoid:Glucocorticoid Ratio | 1,23 | 0.55 | 0.464 | 0.023 | 1,23 | 1.35 | 0.257 | 0.053 | **1,23** | **12.17** | **0.002** ** | **0.336** |
| SMAD specific E3 ubiquitin-ligase 3 | 1,23 | 0.00 | 0.986 | 0.000 | **1,23** | **4.35** | **0.048*** | **0.153** | **1,23** | **4.35** | **0.048*** | **0.153** |

| **Two-way interactions:** | **Age and sex** | | | | **Age and stress reactivity** | | | | **Sex and stress reactivity** | | | |
|---|---|---|---|---|---|---|---|---|---|---|---|---|
| | ***df*** | **F** | ***p*** | **$\eta_p2$** | ***df*** | **F** | ***p*** | **$\eta_p2$** | ***df*** | **F** | ***p*** | **$\eta_p2$** |
| Heart Telomere Length | 1,23 | 1.86 | 0.186 | 0.075 | **1,23** | **10.71** | **0.003** ** | **0.318** | 1,23 | 0.55 | 0.465 | 0.023 |
| Brain Telomere Length | 1,24 | 0.20 | 0.656 | 0.008 | 1,24 | 0.06 | 0.805 | 0.003 | 1,24 | 0.91 | 0.349 | 0.037 |
| Corticotropin-releasing factor | 1,23 | 2.04 | 0.166 | 0.078 | 1,23 | 1.49 | 0.234 | 0.059 | 1,23 | 0.21 | 0.648 | 0.009 |
| Glucocorticoid receptor alpha | 1,23 | 0.32 | 0.577 | 0.013 | 1,23 | 0.09 | 0.768 | 0.004 | 1,23 | 0.68 | 0.417 | 0.028 |
| Mineralocorticoid receptor | 1,23 | 2.22 | 0.149 | 0.085 | 1,23 | 0.11 | 0.742 | 0.005 | 1,23 | 3.20 | 0.086 | 0.118 |
| Mineralocorticoid:Glucocorticoid Ratio | 1,23 | 0.62 | 0.438 | 0.025 | 1,23 | 0.01 | 0.906 | 0.001 | 1,23 | 0.74 | 0.398 | 0.030 |
| SMAD specific E3 ubiquitin-ligase 3 | 1,23 | 0.87 | 0.359 | 0.035 | 1,23 | 3.51 | 0.073 | 0.128 | 1,23 | 0.15 | 0.699 | 0.006 |

| **Three-way interaction:** | **Age, sex and stress reactivity** | | | |
|---|---|---|---|---|
| _ | ***df*** | **F** | ***p*** | **$\eta_p2$** |
| Heart Telomere Length | 1,23 | 0.01 | 0.916 | 0.000 |
| Brain Telomere Length | 1,24 | 2.23 | 0.149 | 0.085 |
| Corticotropin-releasing factor | 1,23 | 0.09 | 0.770 | 0.004 |
| Glucocorticoid receptor alpha | 1,23 | 2.43 | 0.132 | 0.092 |
| Mineralocorticoid receptor | 1,23 | 0.00 | 0.982 | 0.000 |
| Mineralocorticoid:Glucocorticoid Ratio | 1,23 | 2.91 | 0.101 | 0.108 |
| SMAD specific E3 ubiquitin-ligase 3 | 1,23 | 3.53 | 0.072 | 0.128 |

*Significant effects are boldfaced.*

### Supplementary table 3. List of primers used for qPCR analysis

| Gene name (purpose) | Primer name | Accession number | Sequence (5' to 3') | Product size (bp) | Efficiency (%) | Source |
|---|---|---|---|---|---|---|
| **Telomere** | Tel1 | N/A | GGTTTTTGAGGGTGAGGGTGAGGGTGAGGGTGAGGGT | 76 | N/A | https://doi.org/10.1093/nar/30.10.e47 |
| | Tel2 | | TCCCGACTATCCCTATCCCTATCCCTATCCCTATCCCTA | | | |
| **cFos (single copy gene)** | cFos-forward | NM_205569.1 | CAGCTCCACCACAGTGAAGA | 176 | 79 | https://doi.org/10.1371/journal.pone.0086176 |
| | cFos-reverse | | GCTCCAGGTCAGTGTTAGCC | | | |
| **corticotropin-releasing factor (HPI axis)** | crf-zf_qPCR-forward | NM_001007379.1 | CGAGACATCCCAGTATCCAAAAAG | 60 | 111 | https://doi.org/10.1371/journal.pone.0175420.t001 |
| | crf-zf_qPCR-reverse | | TCCAACAGACGCTGCGTTAA | | | |
| **mineralocorticoid receptor (HPI axis)** | mr-zf_qPCR-forward | NM_001100403 | CTTCCAGGTTTCCGCAGTCTAC | 75 | 105 | https://doi.org/10.1371/journal.pone.0175420.t003 |
| | mr-zf_qPCR-reverse | | GGAGGAGAGACACATCCAGGAAT | | | |
| **glucocorticoid receptor α (HPI axis)** | gra-zf_qPCR-forward | NM_001020711.3 | ACTCCATGCACGACTTGGTG | 90 | 92 | https://doi.org/10.1371/journal.pone.0175420.t004 |
| | gra-zf_qPCR-reverse | | GCATTTCGGGAAACTCCACG | | | |
| **SMAD Specific E3 Ubiquitin Protein Ligase** | smurf2-zf_qPCR_forward | NM_001114426.1 | TCAGCCTGGATAAGGGTCAAGG | 100 | 105 | Own produced |
| | smurf2-zf_qPCR_reverse | | TCACCAAGTTCTTAGCGCACAG | | | |
| **ribosomal protein L13a (housekeeping)** | Rpl-F | NM_212784.1 | TCTGGAGGACTGTAAGAGGTATGC | 148 | 80 | https://doi.org/10.3389/fnbeh.2015.00271 |
| | Rpl-R | | AGACGCACAATCTTGAGAGCAG | | | |
| **actin, beta 2 (housekeeping)** | actin-F | NM_181601.5 | CGAGCTGTCTTCCCATCCA | 86 | 95 | https://doi.org/10.3389/fnbeh.2015.00271 |
| | actin-R | | TCACCAACGTAGCTGTCTTTCTG | | | |
